# Ultrasonically Powered Neuromodulation Platform Intended for Spatially and Directionally Selective Electrical Stimulation

**DOI:** 10.1101/2025.05.07.652395

**Authors:** Konstantina Kolovou Kouri, Lukas Holzapfel, Danny Ferreira Dias, Luis Meixner, Wouter A. Serdijn, Vasiliki Giagka

**Affiliations:** Bioelectronics Section, Department of Microelectronics, Faculty of Electrical Engineering, Mathematics and Computer Science, Delft University of Technology, Mekelweg 4, 2628 CD, Delft, The Netherlands; Technologies for Bioelectronics Group, Department of System Integration and Interconnection Technologies, Fraunhofer Institute for Reliability and Microintegration IZM, Gustav-Meyer-Allee 25, 13355, Berlin, Germany; Neuroscience Department, Erasmus Medical Center, Dr. Molewaterplein 40, 3015 GD Rotterdam, The Netherlands

**Keywords:** neuromodulation, selectivity, ultrasound, wireless power transfer, charge balancing

## Abstract

Vagus nerve stimulation (VNS), a well-established application of electrical neuromodulation, has been effective in treating disorders such as epilepsy and depression. Recent neuroscientific advancements have identified further potential in VNS, calling for technical advancements to explore these possibilities. To address this, we developed a neuromodulation platform that can perform both neural activity excitation and inhibition, to further explore spatially and directionally selective stimulation. The platform can be powered wirelessly through ultrasound and includes a charge balancing mechanism to enhance stimulation safety. The platform performs biphasic and monophasic voltage-controlled stimulation, using 6 electrodes for excitation and 2 for inhibition. The pulse width can be varied between 0 and 1500 µs, and the pulse repetition rate between 1 Hz and 1 kHz for excitation and 1 kHz to 50 kHz for inhibition. The implemented charge balancing mechanism uses a digital PID controller to reduce the voltage offset at the stimulation site to values as low as 2 mV. The introduced neuromodulation platform provides a versatile experimental tool intended for, but not limited to, VNS in pre-clinical and clinical settings. It is designed using only commercially available components and is easily reproducible, allowing to explore the possibilities of VNS while identifying the necessary components and processes for developing a miniaturized version.

## Introduction

Vagus nerve stimulation (VNS) has demonstrated great potential in treating a number of neurological and psychiatric disorders, leading to its gradual FDA approval for the treatment of epilepsy, treatment-resistant depression, and post-stroke motor rehabilitation since 1997 (1–3). Nonetheless, the full potential of the therapeutic effects of VNS is still being identified, with promising recent advances in the treatment of autoimmune diseases such as rheumatoid arthritis and Crohn’s disease (4), chronic pain (5) and cardiovascular disease (6), due to its anti-inflammatory effects.

Despite its therapeutic potential, the common practice of stimulating the whole VN and thus the lack of target selectivity during VNS is a cause of unwanted side effects, including bradycardia, cough, hoarseness, and headache (7–9). Recent studies have shown that such side effects can be eliminated by targeting only the specific area of the nerve that includes the fibres of interest, thus offering spatially selective VNS (10, 11). This can be achieved by using multi-contact cuff electrodes, where the contacts can be individually chosen around the circumference of the VN (12, 13). Optimizing VNS through spatial selectivity can give the possibility to better understand the relation to and target specific innervated areas inside the body (14).

The selectivity of VNS when targeting efferent organs can be further enhanced by blocking the activation of the afferent pathways when stimulating the VN and, as a result, only activating the efferent ones. This can be achieved through the application of pulses in the kilohertz domain (kilohertz-frequency alternating current -KHFAC), which has previously been demonstrated to achieve a safe and quickly reversible blocking of the nerve activity (15). Inhibiting the activation of the afferent pathways through KHFAC leads to a temporary virtual vagotomy (transection of the VN) and, thus, directional stimulation selectivity, offering many advantages compared to the equivalent irreversible procedure (16). Given the need to continuously explore the treatment possibilities that VNS has to offer, we developed a neuromodulation platform suitable for investigating the potential of VNS in pre-clinical and clinical settings. The platform is built entirely of commercially available components, making it accessible and reproducible by other research labs to serve as an experimental tool in a variety of applications (17). It is intended for both neural activity excitation and inhibition and can be powered wirelessly through ultrasound (US), using the same signal for powering and stimulation.

Our proposed system has an output stage that supports a connection to 8 electrodes, two for the purpose of neural activity inhibition and six for excitation. By providing the capability to inhibit the propagation of action potentials along the nerve, while simultaneously exciting the nerve at a different location, we can achieve directional selectivity. Furthermore, each of the six excitatory electrodes can be individually selected and controlled, allowing the manipulation of the generated electric field and thus adding a degree of spatial selectivity of the activated area. The system delivers monophasic or biphasic electrical stimulation.

Due to the non-linear behaviour of the electrode-tissue interface (ETI) (18–20) and possible mismatches in the stimulation pulse phases due to hardware limitations and electronic component tolerances, charge can accumulate at the ETI, leading to the buildup of a residual DC voltage, which might cause irreversible electrochemical reactions, harming either the tissue or the electrode (21, 22). A charge balancing scheme using a Proportional-Integral-Derivative (PID) controller (23, 24) is implemented here to reduce the excess charge appearing at the stimulator output.

Aiming towards an eventual miniaturization of the entire system, as envisioned in (25) and seen in figure 1, the platform is powered wirelessly through US. To increase the system’s power efficiency, thereby reducing power dissipation and thus heat generation, the harvested signal used to power the system is also used for neural stimulation (26, 27). The rectified sinusoid from the energy harvester is directly forwarded to the system’s output stage and the stimulating electrodes, eliminating the need for an additional voltage or current source. Furthermore, the neuromodulation platform can be powered through a USB connection, or a battery for portability. The user can communicate with the platform through a graphical user interface (GUI) on a laptop or PC.

**Fig. 1.**
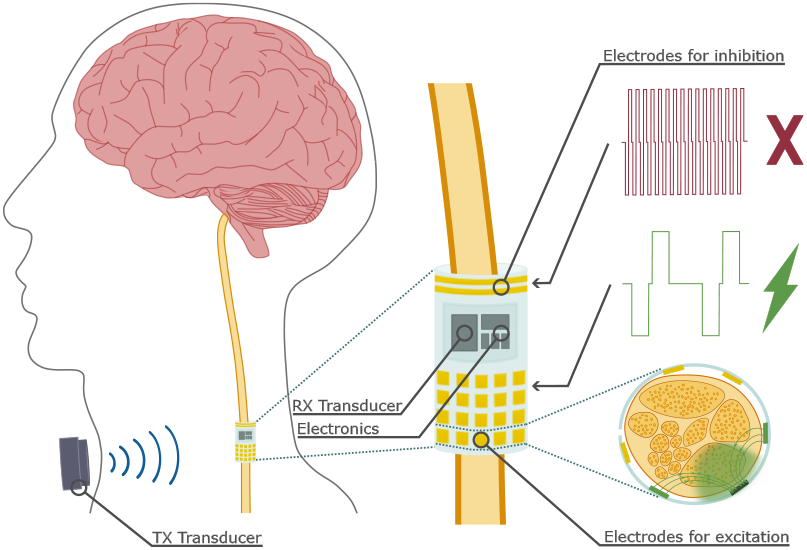
Concept of wireless battery-free neurostimulator implant. The miniaturised implant is powered wirelessly through ultrasound and uses the same harvested ultrasonic signal, converted into the electrical domain, for both powering and neural stimulation. The combination of direction-specific stimulation through KHFAC block and area-specific excitation through individually addressable electrodes allows for selective stimulation of the nerve’s fascicles and fibres, depending on the targeted area. A benchtop neurostimulator platform is developed, which can be used to identify the most relevant parameters for VNS as well as the most effective and energy-efficient design for a miniaturised battery-free neural implant.

## Materials and Methods

A block diagram of the proposed neuromodulation platform is presented in figure 2a. The system consists of five main blocks. Acoustic energy is harvested, converted into the electrical domain and rectified within the Energy Harvesting block. This rectified signal can subsequently be used to power the platform and for stimulation at the output stage. An alternative power source is provided through a battery or a USB connection (also allowing for data transfer) and is part of the Power Supply & Data Transfer block. The Power Management block is responsible for all necessary power conversions to supply the system’s components, while all logic operations are performed within the Control Unit block. Finally, the Output Stage enables the connection to the stimulation site. A detailed description of all individual blocks and components follows.

**Fig. 2.**
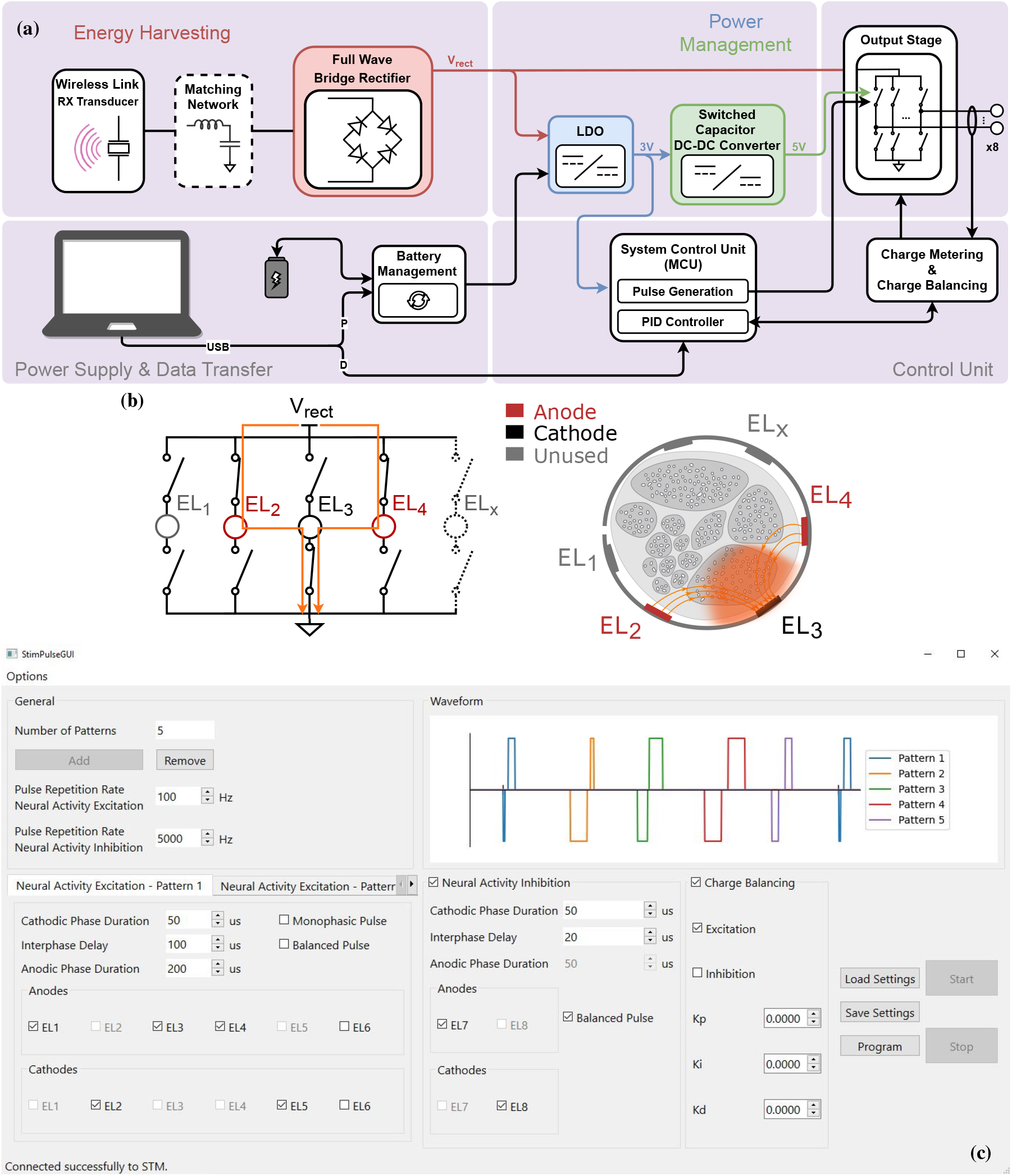
a) System-level block diagram of the proposed neuromodulation platform and its electrical functionalities. The platform can be powered wirelessly, with the harvested US signal being used to generate regulated supplies to power the various blocks of the system as well as directly for neural modulation. For testing purposes and for the identification of all relevant stimulation parameters, the US signal can be emulated through a function generator, while the rest of the system can be powered through a battery or a USB connection. The user can communicate with the platform through a GUI on a laptop or PC through the same USB connection. b) The possibility to customize the configuration of the electrodes used for neural activity excitation allows for manipulation of the generated electric field, leading to spatial selectivity of the activated area. Here, electrodes 2 and 4 are chosen as anodes, while electrode 3 is the cathode, concentrating the electric field and stimulation area around electrode 3. c) Screenshot of the GUI used to configure the stimulation parameters.

### A. Neurostimulator Circuit Hardware

The neuromodulation platform can be powered through multiple sources. A USB connection to a laptop or PC provides a constant 5V supply to the system. When forwarded to the battery charge-management IC (MAX8606), it can recharge a 3.7 V lithium polymer battery with 2000 mAh current capacity (RS Pro). Disconnecting from the USB supply and using the battery pack to provide the necessary power allows for a portable autonomous system that does not require any external connections. Finally, the platform can be powered wirelessly through US. The harvested acoustic energy is converted into the electrical domain through a receiving US transducer (RX). Depending on the transducer used as RX, a matching circuit can be included to maximize power transfer. The harvested signal is subsequently rectified, with four Schottky diodes (RB751CM-40) connected in a bridge circuit configuration providing full-wave rectification.

The rectified voltage is forwarded to the upcoming power management blocks as well as the output stage of the system. A low-dropout (LDO) voltage regulator (TPS79730) regulates the rectified US signal to a constant 3 V to power a microcontroller unit (MCU), the STM32L562. Following the LDO, a step-up charge-pump DC-DC converter (MAX619) provides a regulated 5 V to operate four single-pole, single-throw (SPST) analog switches (TMUX1112), each with four independently selectable channels, and three quad-channel operational amplifiers (OpAmps) (TLV9044). Both the TMUX1112 and the TLV9044 offer rail-to-rail input and output.

The same signal used for powering the platform is forwarded to the system’s output for stimulation, thereby achieving voltage-controlled stimulation. This approach bypasses the requirement for an additional voltage or current source that is typically used in electrical stimulators and avoids unnecessary power conversions, thus implementing a more energy-efficient version of the system. When using the battery pack or the USB connection to supply the power, which is also useful for debugging and testing purposes, the system is decoupled from the wireless power supply. In this case, the US signal needed for stimulation can be emulated through a function generator.

The system’s output stage supports a connection to eight electrodes, two for the purpose of neural activity inhibition and six for excitation. A choice can be made for either monophasic or biphasic stimulation. Monophasic stimulation offers the advantage of lower stimulation thresholds and higher spatial selectivity (28); however, it compromises safety, which is why biphasic stimulation is preferred in recent stimulation protocols. Given the need for a biphasic pulse for neural stimulation, an H-bridge configuration at the output stage, using the TMUX1112 analog switches, allows for stimulation with both polarities while using one single power line - in this case, the full-wave rectified signal from the US transducer. This way, the two stimulation phases, cathodic and anodic, are generated by changing the direction of the current that flows through the tissue. An example of the H-bridge switches’ configuration for one phase is depicted in figure 2b. In this scenario, electrodes 2 and 4 are chosen as anodes, while electrode 3 is chosen as a cathode. By closing the upper H-bridge switches of electrodes 2 and 4 and the lower H-bridge switch of electrode 3, current can flow from electrodes 2 and 4, through the tissue, to electrode 3. In the next phase, to reverse the current flow, only the lower H-bridge switches of electrodes 2 and 4, and the upper H-bridge switch of electrode 3 will be closed (not depicted here).

The purpose of a balanced biphasic pulse is to inject and remove the same amount of charge needed to stimulate the nerves so as not to cause excess charge buildup at the ETI that might harm the tissue and damage the electrode (21). For this reason, it has been customary in the design of neural stimulators to balance the two stimulation phases as accurately as possible (29), while also including a coupling capacitor to prevent charge buildup (30). However, even a perfectly balanced biphasic pulse will not always guarantee the absence of harmful electrochemical reactions, given the non-linear nature of the ETI. To mitigate this, stimulator designs started introducing features that aim at actively reducing any charge buildup that leads to a voltage offset detected by the electronic system at its output after the end of a stimulation pulse (31).

A good practice to minimize the charge buildup is by eliminating the offset at the electrodes connected to the output terminals of the system by means of negative feedback (32). Ideally, any electrode voltage offset should be minimized versus a non-polarizable reference electrode with a stable potential at the tissue side. Here, in the absence of in vivo non-polarizable reference electrodes, one of the electrodes of our system is defined as reference, placed at the tissue and gets assigned a stable potential, against which the potential at every stimulating electrode is compared. By bringing all other electrode potentials to the potential of the reference electrode, all inter-electrode voltages are close to zero, thus reaching a zero net charge at the stimulation site. To achieve this, a circuit is needed to monitor the voltage at each electrode.

A second-order active low-pass (LP) filter in a Sallen–Key topology is placed behind each electrode to monitor the electrode’s voltage with respect to ground and to forward this voltage to the MCU for further processing. Each filter consists of one unity-gain OpAmp, part of the TLV9044 quad-channel OpAmps, and two RC networks. By choosing the RC network values at *R* = 1 MΩ and *C* = 15 nF, the cut-off frequency *f*_c_ of the LP filter is set at approximately 10 Hz, since

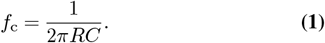

Given the low *f*_c_ of the filter, as well as its order, which leads to an attenuation by 40 dB*/*decade, this essentially allows for a DC measurement of the electrode voltage.

As previously mentioned, to achieve a zero potential difference between the stimulating electrodes and, thus, a zero average differential output voltage at the stimulation site, the resting potential of all electrodes needs to be at the same level. For this, the measured electrode voltage is compared to a reference voltage *V*_ref_. This *V*_ref_ is set halfway between the supply voltage and ground, as the common-mode output voltage, and assigned to the reference electrode. When powering with US, the supply voltage corresponds to the voltage *V*_rect_ that is generated by harvesting the acoustic energy, converting it to the electrical domain and rectifying the resulting sinusoidal signal. Consequently, *V*_ref_ is set at *V*_rect_*/*2.

To ensure a reliable *V*_ref_, an accurate reading of the supply voltage *V*_rect_ is needed. An active LP filter with a cut-off frequency of *f*_c_ = 10 Hz provides a precise value of *V*_rect_. A voltage divider contributes to the LP filter behaviour while producing the necessary *V*_rect_*/*2 to be used as *V*_ref_. Finally, this value is forwarded to the MCU, where the necessary actions are taken to obtain a zero net charge between all electrodes of the stimulation site.

Every electrode of the output stage is controlled by one side of the H-bridge, giving the possibility to set each electrode individually as anode or cathode. This feature makes spatially selective stimulation feasible, as the selection of more than one electrode as an anode can help manipulate the generated electric field and concentrate it to a specific region within the nerve, as seen in figure 2b.

The main specifications of the complete system are summarized in table 1.

**Table 1.** Neuromodulation platform specifications overview.

| Parameter | Value |
| --- | --- |
| Nr. of electrodes A<sup>a</sup> | 6 |
| Nr. of electrodes I<sup>a</sup> | 2 |
| Pulse pattern | Mono-/biphasic symmetric/asymmetric |
| Pulse polarity | - (cathodic) or + (anodic) first |
| Pulse width | 0 - 1500 $\mu\text{s}$ (in 1 $\mu\text{s}$ steps) |
| Interphase delay | 5 - 200 $\mu\text{s}$ (in 1 $\mu\text{s}$ steps) |
| PRR<sup>b</sup> A<sup>a</sup> | 1 Hz - 1 kHz |
| PRR<sup>b</sup> I<sup>a</sup> | 1 kHz - 50 kHz |
| Supply | Wirelessly through US<sup>c</sup><br>LiPo<sup>d</sup> 3.7 V 2 mAh cells (rechargeable)<br>5 V through USB |

<sup>a</sup>A and I denote Activation and Inhibition, respectively;
<sup>b</sup>Pulse Repetition Rate;
<sup>c</sup>Ultrasound;
<sup>d</sup>Lithium Polymer.

### B. Graphical User Interface

A Graphical User Interface (GUI) is used to define the stimulation settings, as seen in figure 2c. The visual part of the GUI is designed using the PySide6 Designer. Using the designer’s interface, all necessary windows, text fields, and buttons are selected and arranged in the desired pattern. From this design, the “.ui” files are generated and subsequently imported into Python, where all windows, buttons, and fields are linked to their functions, allowing the GUI to operate as intended. The library of the Protocol Buffers (`protobuf`), a mechanism for serializing structured data, is used so that the GUI can communicate with the MCU.

Through the GUI, the parameters for the neural activity excitation and inhibition can be set. Each of the stimulating electrodes can be individually selected and defined as either anode or cathode. At least one of the electrodes needs to be selected as anode and one as cathode to ensure current flow between the electrodes. An electrode cannot have both anode and cathode status to avoid shorting at the stimulation site. To increase flexibility in the stimulation pulse sequence, up to five different biphasic pulse patterns can be programmed within one time period, as seen on the waveform visualization in figure 2c. These five stimulation patterns can be used only for the purpose of neural activity excitation, and their parameters can be configured for each pattern individually. Among those configurable parameters are the pulse repetition rate (PRR), i.e., the time frame in which all patterns are repeated, the cathodic and anodic phase duration and the interphase delay duration, and finally, the selection of the electrodes and their allocation as anodes or cathodes. The same parameters can be configured for inhibitory patterns, which can be activated and deactivated depending on the stimulation needs.

### C. Firmware

The STM32L562 microcontroller (µC) is programmed in C language using the STM32CubeIDE 1.9.0.

#### C.1. Stimulation Pulse Generation

The timers of the µC are used to provide the control signals for stimulation. By using the complementary pulse width modulation (PWM) function of the timers, a biphasic signal can be generated using the regular channel output (`TIMx_CHy`) for the first phase and the complementary channel output (`TIMx_CHyN`) for the second phase of the biphasic pulse. A dead time (DT) can be inserted between the two outputs, acting as an interphase delay and ensuring that the two outputs are not active simultaneously. The advanced-control timers `TIM1` and `TIM8` are used for the excitatory and the general-purpose timer `TIM15` for the inhibitory signals. Timers `TIM1` and `TIM8` each have three channels, all with a regular and a complementary output, while timer `TIM15` has two channels, out of which only one has a complementary output.

To achieve the necessary flexibility for spatially selective stimulation, each µC channel controls only one side of the H-bridge, i.e., one electrode at the output stage. Here, the regular channel output `TIMx_CHy` controls the upper switch and the complementary channel output `TIMx_CHyN` controls the lower switch. Therefore, a total of six µC channels (`TIM1` and `TIM8`) can control a maximum of six electrodes used for excitation.

Responsible for triggering, and thus the synchronised start, of the timers `TIM1` and `TIM8` is the general-purpose timer `TIM4`. Whenever `TIM1` or `TIM8` are triggered by `TIM4`, the first phase of the biphasic pulse commences (`TIMx_CHy` is “high”), together with the incrementation of each timer’s own counter. The first phase ends when reaching the value of the CCR register, at which point, the timer changes polarity (`TIMx_CHyN` is “high”). A dead time can be assigned as an interphase delay between the two phases of the pulse. The second phase commences right after the dead time, with its end being indicated by the interrupt service routine (ISR) of `TIM1`, which is called at the moment the timer counter overflows. The length of the second pulse is defined as

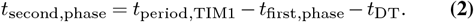

In a similar manner, the general-purpose timer `TIM3` triggers the start of the biphasic pulse for the inhibition, conducted by `TIM15`. The ISR of `TIM15` ends the pulse. As there are only two electrodes for inhibition, their control can be achieved by a single H-bridge. Thus, the architecture that is used for excitation, enabling the combination of different electrodes, is no longer needed. Instead, a single µC channel, connected to both electrodes, is sufficient to perform the switching. For this, one of the two µC channels of `TIM15` is needed, the one with the complementary output.

Whenever the electrodes for inhibition are not stimulating, they are left floating to prevent any current flow between them, which is achieved by leaving all channels open. When the electrodes for excitation are not stimulating, they are grounded to remove any remaining charge built up. This is achieved by connecting all complementary channels `TIMx_CHyN`, and thus the lower part of the H-bridge, to ground. The latter approach cannot be implemented for the inhibitory electrodes, as the switches of the H-bridge used for inhibition cannot be individually controlled. The timings and dependencies of the timers and the resulting H-bridge control are illustrated in figure 3a.

**Fig. 3.**
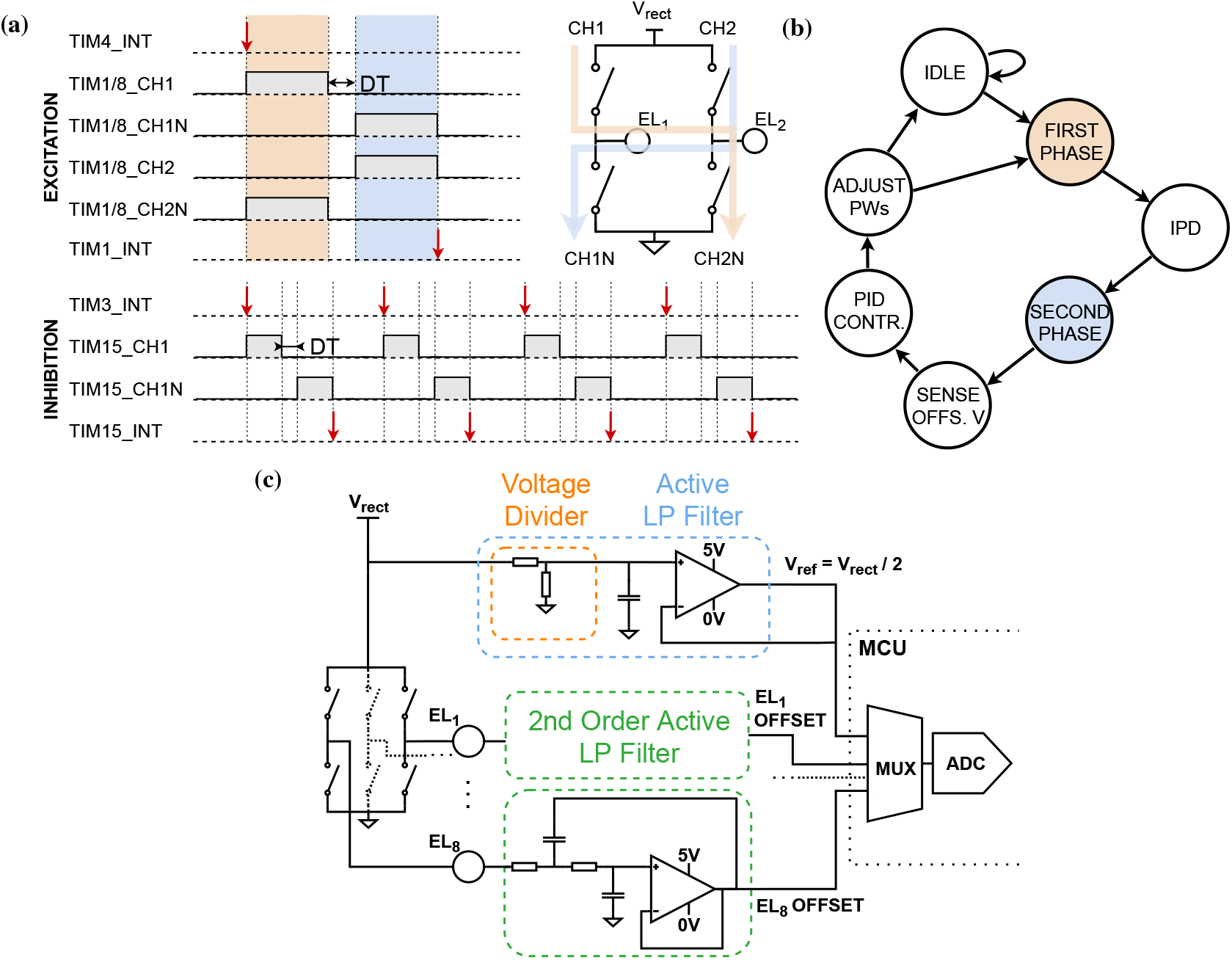
a) Time diagram of the biphasic stimulation pulse generation within the STM32L562 µC. The interrupt service routine (ISR) of the general-purpose timer TIM4 triggers the start of the advanced-control timers TIM1 and TIM8 that control the excitatory electrodes. The first stimulation phase (orange) is followed by the interphase delay, set by the dead time register of the microcontroller, and by the second phase (blue), thus controlling the two sides of the H-bridge at the output stage. The biphasic pulse ends with the ISR of TIM1, which is called at the overflow of the timer counter. The same routine is followed for the timers that control the inhibitory electrodes. b) Finite state machine (FSM) representation of a biphasic pulse sequence and the control loop actions for charge balancing. c) Detail of the output stage and the circuitry used to accurately measure the values needed for charge balancing. The electrode voltage at each electrode is measured with a second-order active low-pass filter and is compared to a reference voltage *V*_ref_ for the purpose of achieving a zero potential difference between any stimulating electrode pairs. *V*_ref_ is equal to *V*_rect_*/*2, obtained by passing *V*_rect_ through a voltage divider and an active low-pass filter.

#### C.2. PID Controller for Charge Balancing

A charge balancing scheme is implemented to remove as much excess charge as possible from the ETI after a stimulation cycle, which might cause irreversible reactions to either the tissue or the electrode. The particularity of this design is that the signal used for neural stimulation is directly derived from the signal used for powering the device and, therefore, its amplitude depends on the intensity of the harvested US signal and cannot be changed during stimulation. As a result, in order to modify the amount of charge injected into the tissue, only the duration of the pulses can be adapted accordingly.

This pulse width adaptation is conducted by a digital PID controller running on the µC, which regulates the pulse width of both phases of every upcoming biphasic pulse to avoid excess charge buildup. Generally, a control loop is used to bring a specified physical variable, the measured process variable (PV), to a desired value (set-point, SP) and maintain it at that, regardless of disturbances. In order to fulfil the control task, the instantaneous value of the PV must be measured and compared with the SP value. Any deviations from the desired SP are re-adjusted accordingly through the controller’s output. Here, the desired SP is *V*_rect_*/*2, as explained in section A. The PV is the voltage measured at each electrode of the output stage. This voltage is sampled during stimulation, acting as feedback to be compared with the desired SP. Before any of the signals can be processed by the µC, the measured PV and SP must first be digitised using an analog-to-digital converter (ADC). With the ADC clock frequency at 20 MHz and the sampling time set at 640.5 clock cycles (total conversion time = sampling time + 12.5 ADC clock cycles), the ADC can sample every 32.65 µs. The necessary ADC conversions amount to 9, one for the voltage of each of the PVs plus one for the SP, *V*_rect_*/*2. Given this number and the ADC sampling time, the ADC needs a total of 293.85 µs for all 9 conversions, which corresponds to a sampling rate of approximately 3.4 kHz. Choosing this relatively low sampling rate for the ADC is beneficial in terms of energy efficiency. Given that the measured PV can be considered a DC measurement (because it is LP-filtered at 10 Hz), minimizing the effective sampling rate of the ADC to 3.4 kHz reduces the overall power consumption.

Finally, the output signal is calculated by the digital controller. This output dictates the amount of time by which the cathodic and anodic phases need to be increased or decreased to mitigate any leftover potential difference between the electrodes. Since the total pulse width of one biphasic pulse (cathodic phase + interphase delay + anodic phase) is fixed due to the chosen method of generating these pulses (see equation 2), for charge balancing, only the ratio between the phases of each pulse can be adapted. The controller measures the PV after every stimulation pulse, compares it to the SP and computes the pulse width adaptation for the next pulse.

The finite state machine in figure 3b illustrates the process of the biphasic pulse generation and the charge balancing method using the PID control loop. Starting from the IDLE state, the first phase of the biphasic pulse commences as soon as the command is sent through the GUI. Based on the chosen electrode configuration, this can be defined as either anodic or cathodic. The first phase is followed by the interphase delay and the second phase of opposite polarity. The value of the sampled electrode voltage is sent as feedback to the PID controller, which outputs the amount of time by which the two phase durations need to be adjusted. If the stimulation sequence is stopped, the system returns to the IDLE state; otherwise, it continues with the generation of the next biphasic pulse.

The digital implementation of the PID controller in the µC requires the standard form of the PID controller to be implemented in a discrete-time fashion. The equation for the discrete-time system is created based on the equation for the continuous system. Through the approximation of the individual terms by using backward finite differences, the discrete implementation of the controller output equation results in

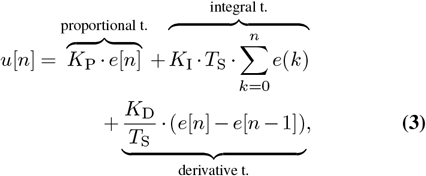

comprised of the proportional, integral and derivative terms, where K_P_, K_I_ and K_D_ denote the proportional, integral and derivative gain, respectively. *e* indicates the error signal that the controller uses to calculate its output, which is the difference between the measured PV and the desired SP. *T*_S_ is the sampling time between two sampling points and *n* indicates the sampling number. The controller output value is then added to or subtracted from the pulse width of the cathodic or anodic phase. The controller is furthermore equipped with an anti-windup mechanism; when the integral term reaches saturation, either at its upper or lower bound, it is clamped to the corresponding predefined limit. As a result, the integral term cannot grow uncontrollably.

The mechanism for charge balancing can be activated or deactivated through the GUI. Furthermore, the GUI allows for the modification of the proportional, integral and derivative gains, adjusting them accordingly to fit the individual needs of the electrodes interfacing with the neuromodulation platform and the stimulation pattern characteristics. Finally, any of the K_P_, K_I_ or K_D_ gains can be set to zero, allowing for a different version of the controller.

### D. System Validation

To validate the functionality and performance of the neuromodulation platform, experiments are conducted in two different configurations. For the first one, the US signal is emulated either in its harvested AC form through a function generator (33622A, Keysight) or in its rectified form through a power supply - either a laboratory power supply (EDU36311A, Keysight) or the available USB connection. This gives a clear overview of the functionality of the individual components of the platform.

For the second configuration, the platform is entirely powered through US. Two commercially available transducers by Precision Acoustics, suitable for immersion use, are used as the transmitter (TX) and receiver (RX). The TX has a diameter of 19 mm and a resonance frequency of 2 MHz, matching the resonance frequency of the RX that has a diameter of 15 mm. Both the TX and the RX are immersed in a tank filled with deionized (DI) water. The water tank is layered with acoustic absorbers (VK-76000, Gampt), to eliminate the unwanted reflections that could influence the results of the measurements.

The output of the neuromodulation platform, as well as all intermediate signals, are displayed on an oscilloscope (MDO34, Tektronix and RTA4004, Rohde&Schwarz). The parameters for a stimulation protocol are set and changed through the GUI on a laptop. Figure 4 illustrates the setup used for all measurements.

**Fig. 4.**
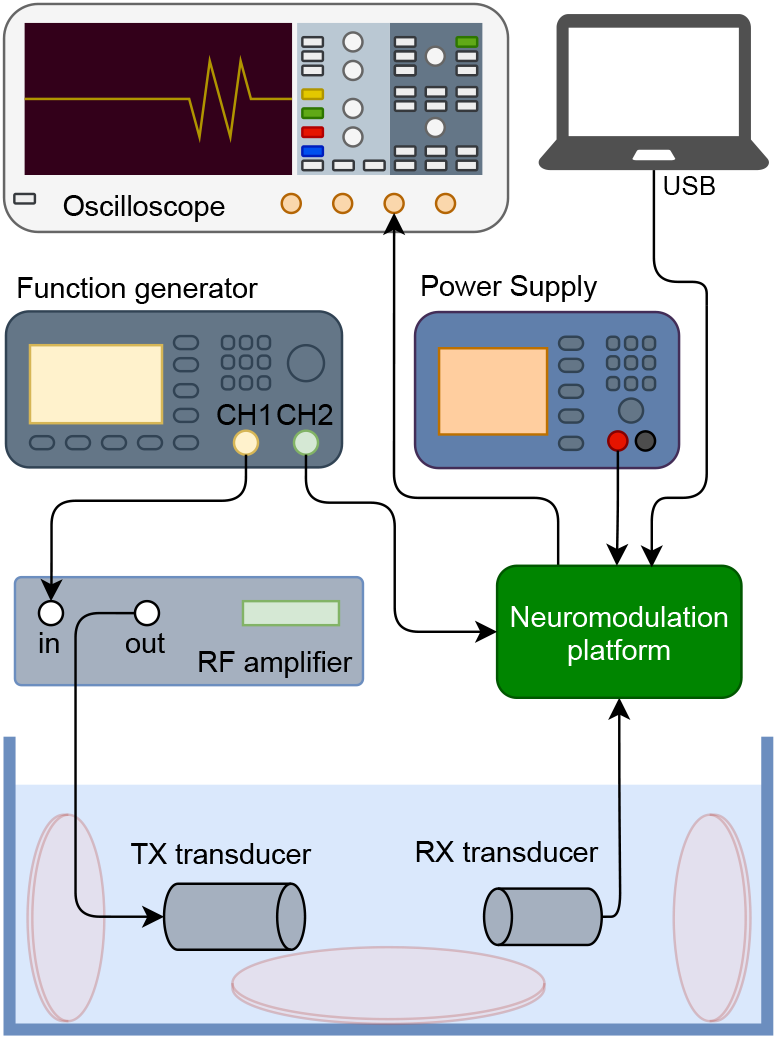
Schematic of the measurement setup for the functionality validation of the neuromodulation platform.

For the initial system validation measurements, the load connected to the output stage is purely resistive. All further measurements are conducted with electrodes submerged in a phosphate-buffered saline (PBS) solution (0.14 M NaCl, 2.7 mM KCl, 10 mM phosphate buffer, with a pH value of 7.4 *±* 0.05), which mimics the physiological environment of the inner body. The electrodes used are the Dorsal Root Ganglion (DRG) leads from Abbott^©^ (Proclaim™ DRG), equipped with 4 electrodes. All 4 electrodes are characterized prior to the validation measurements through electrochemical impedance spectroscopy (EIS), with the graph in figure 5 presenting an EIS measurement between 2 adjacent out of the 4 electrodes.

**Fig. 5.**
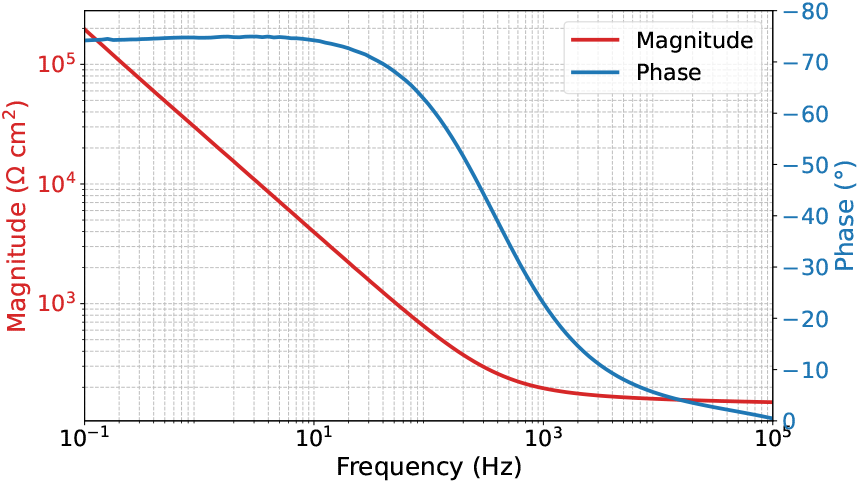
Electrochemical impedance spectroscopy (EIS) of the DRG leads from Abbott^©^ (Proclaim™ DRG).

## Results

### E. Wireless Link Characterization

To successfully power the neuromodulation platform with US, the performance of the wireless US link is assessed. Figure 6a illustrates the TX and RX voltage, *V*_TX,pp_ and *V*_RX,pp_, respectively, along with the rectified voltage *V*_rect_ that is eventually used to power the platform. This measurement is conducted in an open-circuit configuration, reducing the voltage losses to a minimum and providing an indication of the maximum available power that can be transferred in this configuration. Here, for a *V*_RX,pp_ = 10.65 V, we obtain a *V*_rect_ = 9.72 V - given that the diodes of the bridge rectifier and the parasitic capacitances of the RX transducer form a voltage multiplier.

**Fig. 6.**
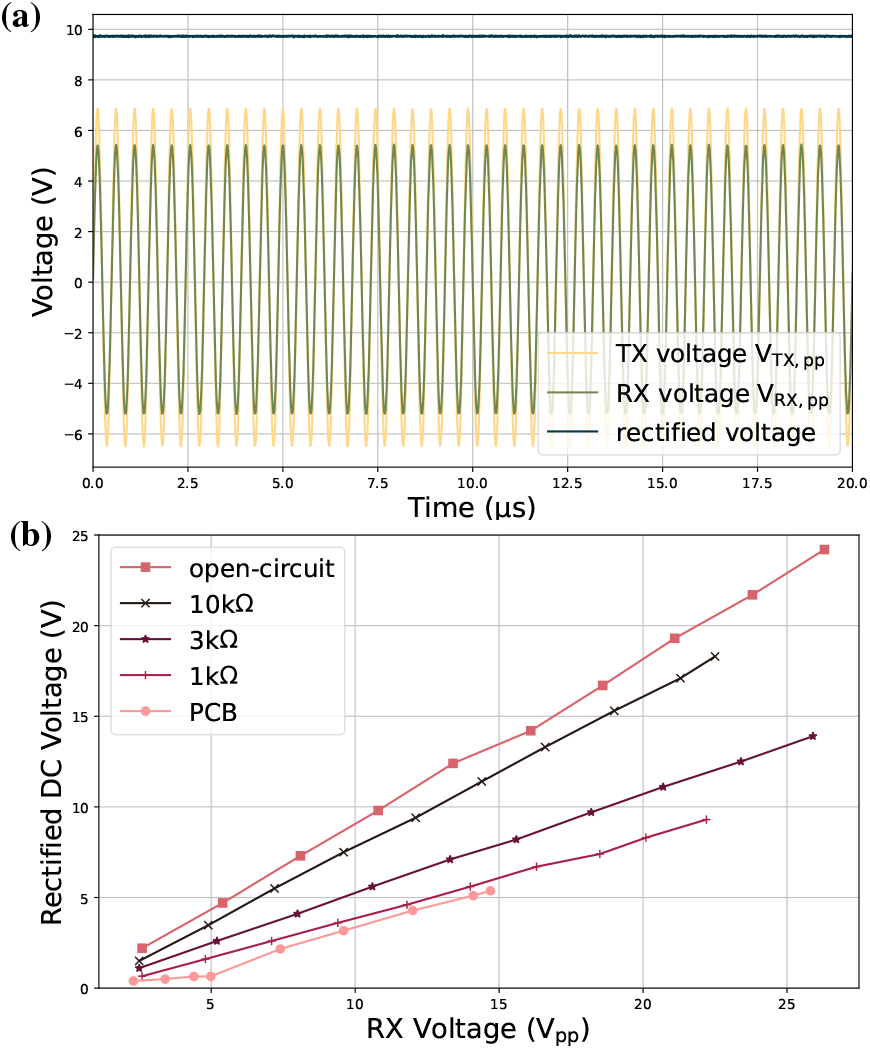
a) Representation of the voltage at the US transmitter TX *V*_TX,pp_, the voltage at the US receiver RX *V*_RX,pp_ and the rectified voltage *V*_rect_ that is used to power the neuromodulation platform, with the measurement conducted in an open-circuit configuration. b) Development of the rectified voltage *V*_rect_ as a function of the voltage at the US receiver RX *V*_RX,pp_ when connecting to different loads.

Furthermore, the obtained rectified voltage *V*_rect_ is observed as a function of the voltage at the US RX *V*_RX,pp_ when connecting it to different loads. *V*_rect_ is measured in an open-circuit configuration, for three different resistive loads, and finally for when the rest of the platform is connected and acts as a load. For the last case, *V*_RX,pp_ is increased only until *V*_rect_ reaches the value of 5.5 V, which is the maximum voltage that can serve as an input to the LDO that follows the rectifier on the platform. Figure 6b depicts the development of *V*_rect_ with an increasing *V*_RX,pp_.

### F. PID Controller Behaviour and Charge Balancing

The PID controller used for charge balancing is parametrized empirically using the DRG leads from Abbott^©^ submerged in a PBS solution. One of the electrodes on the Abbott^©^ DRG leads is connected to the reference voltage *V*_ref_ (here at *V*_rect_*/*2) that is used as the SP. That way, we provide a stable potential within the same environment as the stimulating electrodes, against which we can compare the measured electrode voltage to eliminate any potential difference between all electrodes. The behavior of the controller is analyzed for different proportional, integral and derivative gains, resulting in an optimum combination for the particular application.

Starting with the tuning of the proportional gain K_P_, K_P_ is adjusted while K_I_ and K_D_ are set to zero. K_P_ is gradually increased, resulting in an increasingly aggressive controller. With the response of the control loop becoming more reactive, the deviation of the PV from the desired SP becomes smaller until the system becomes unstable and starts to oscillate. Figure 7a demonstrates this behavior for seven different K_P_ values. K_P_ = 0.8 is chosen to continue the tuning for the integral and derivative gains, as it approaches the desired SP the closest and without oscillations.

**Fig. 7.**
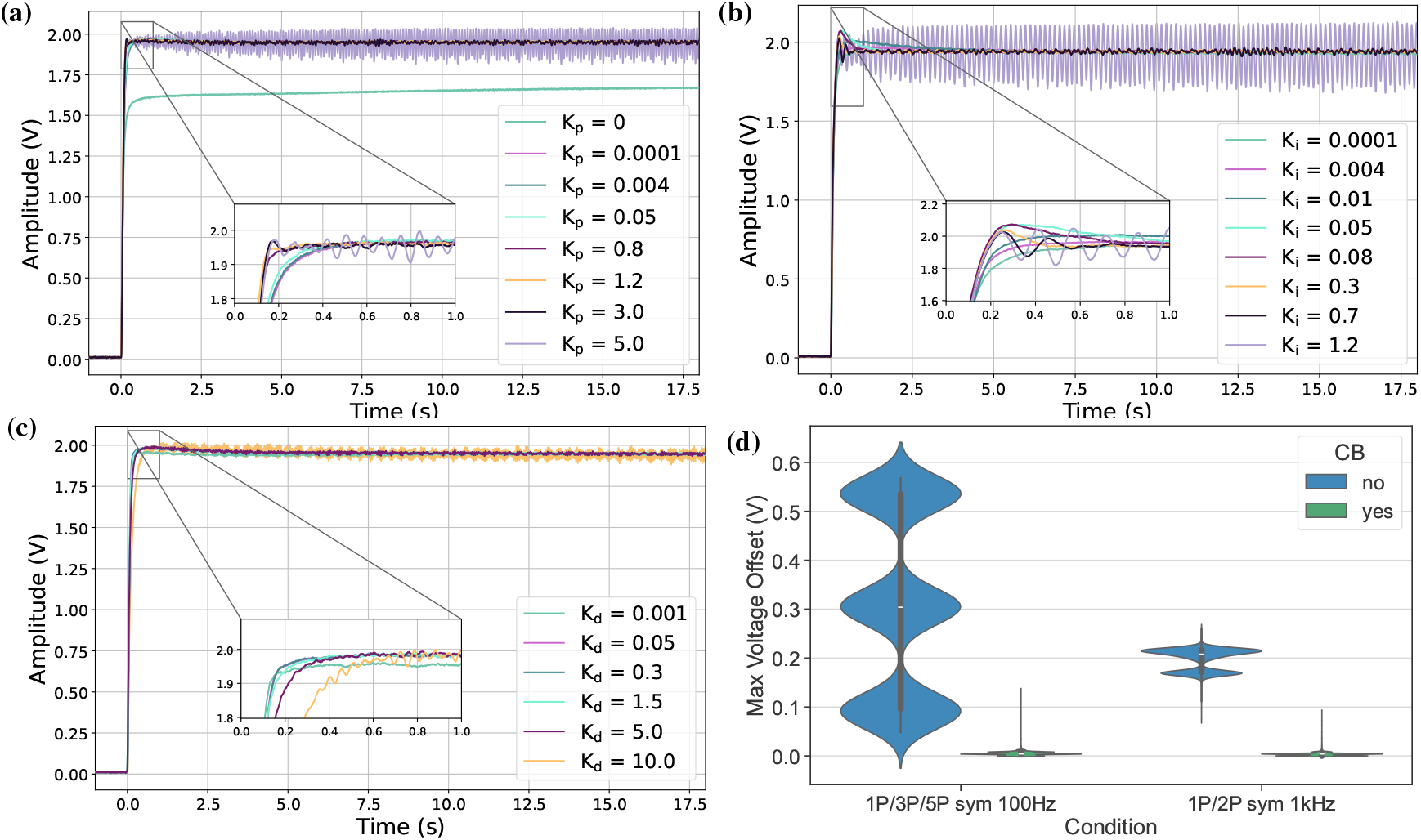
Parametrization of the PID controller used for charge balancing. a) K_P_ is varied over multiple values. For K_P_ = 0.8, the deviation between the process variable (PV) and the desired set-point (SP) decreases without the controller oscillating. b) K_I_ is varied over multiple values. For K_I_ = 0.004, the controller settles faster without unwanted oscillations or instability. c) K_D_ is varied over multiple values. For K_D_ = 0.001 the controller settles slightly faster to the desired SP. d) Maximum potential difference (voltage buildup due to charge accumulation) between stimulating electrodes without and with the charge balancing mechanism activated.

With K_P_ set as 0.8, K_I_ is now varied while K_D_ remains zero. Similarly to the settings for K_P_, as the value for K_I_ increases, so does the aggressiveness of the controller. A too-low value leads to a (possibly too) slow settling of the controller, while a too-high value leads to large oscillations and unstable behavior, as can be seen in figure 7b. Furthermore, a higher than necessary K_I_ value can lead to a large overshoot and potentially the injection of a higher voltage that might harm the stimulated tissue. K_I_ was chosen as 0.004, for which the controller settles quickly to the desired SP without any oscillations or overshoot.

Finally, the same procedure is followed for the derivative gain K_D_. K_D_ is varied over six values, with no significant changes for lower K_D_ values, as can be seen in figure 7c. When increasing the value for K_D_, the controller starts becoming unstable. For this reason, K_D_ is set to 0.001. All gains can be varied as needed depending on the application and type of electrodes that the tuning is intended for.

After tuning the controller to the intended measurement setup, the potential difference between all stimulating electrodes is measured for a variety of settings. The maximum potential difference is calculated and compared between having the charge balancing mechanism deactivated and activated. For this, we varied the PRRs (100 Hz vs. 1 kHz) of the stimulation pulses, as well as the number of patterns employed for every stimulation sequence (1, 3 and 5 patterns (1P/3P/5P) for a PRR of 100 Hz, and 1 and 2 patterns (1P/2P) for a PRR of 1 kHz). Figure 7d demonstrates the effectiveness of the employed charge balancing mechanism, lowering the potential difference for lower PRRs (100 Hz) from approximately 300 mV to 3 mV, and for the higher PRRs (1 kHz) from approximately 200 mV to 2 mV, where all values are mean values.

### G. System Output Performance

#### G.1 Biphasic Pulse and Multichannel Operation

The output of the neuromodulation platform, as well as all intermediate signals, are displayed on an oscilloscope (MDO34, Tektronix and RTA4004, Rohde&Schwarz). The parameters for a stimulation protocol are set and changed through the GUI on a laptop. Figure 4 illustrates the setup used for all measurements. Up to a total of five different patterns can be activated for neural excitation within one stimulation period, allowing for interleaved stimulation. Each pattern can have an independent set of parameters, thus addressing a different group of electrodes while having a different pulse width and inter-phase delay. An illustration of the patterns appears on the GUI, as shown in figure 2c, to aid the user in visualising the programmed stimulation protocol. Furthermore, the output stage of the platform can connect to two electrodes for neural activity inhibition, and the connection can be activated or deactivated at will. The biphasic pulse generation and the flexibility of the multichannel operation are evaluated while powering the platform through the USB connection and emulating the US signal used for stimulation through the 33622A Keysight function generator (signal amplitude of 2 V).

Figure 8a shows the measured voltage at the platform’s output when all 5 patterns are being engaged, with the settings corresponding to the ones displayed in figure 2c. This graph demonstrates the accuracy of the neuromodulation platform in producing the desired patterns and the possibility of engaging up to 5 stimulation patterns within one period of a stimulation protocol. The plot shows the measured voltage when connecting the platform’s output to purely resistive loads of 1 kΩ and 100 kΩ. Even though multiple electrodes can be engaged within one pattern, figure 8a only demonstrates the voltage measured between two electrodes, for ease of comparison with figure 2c.

**Fig. 8.**
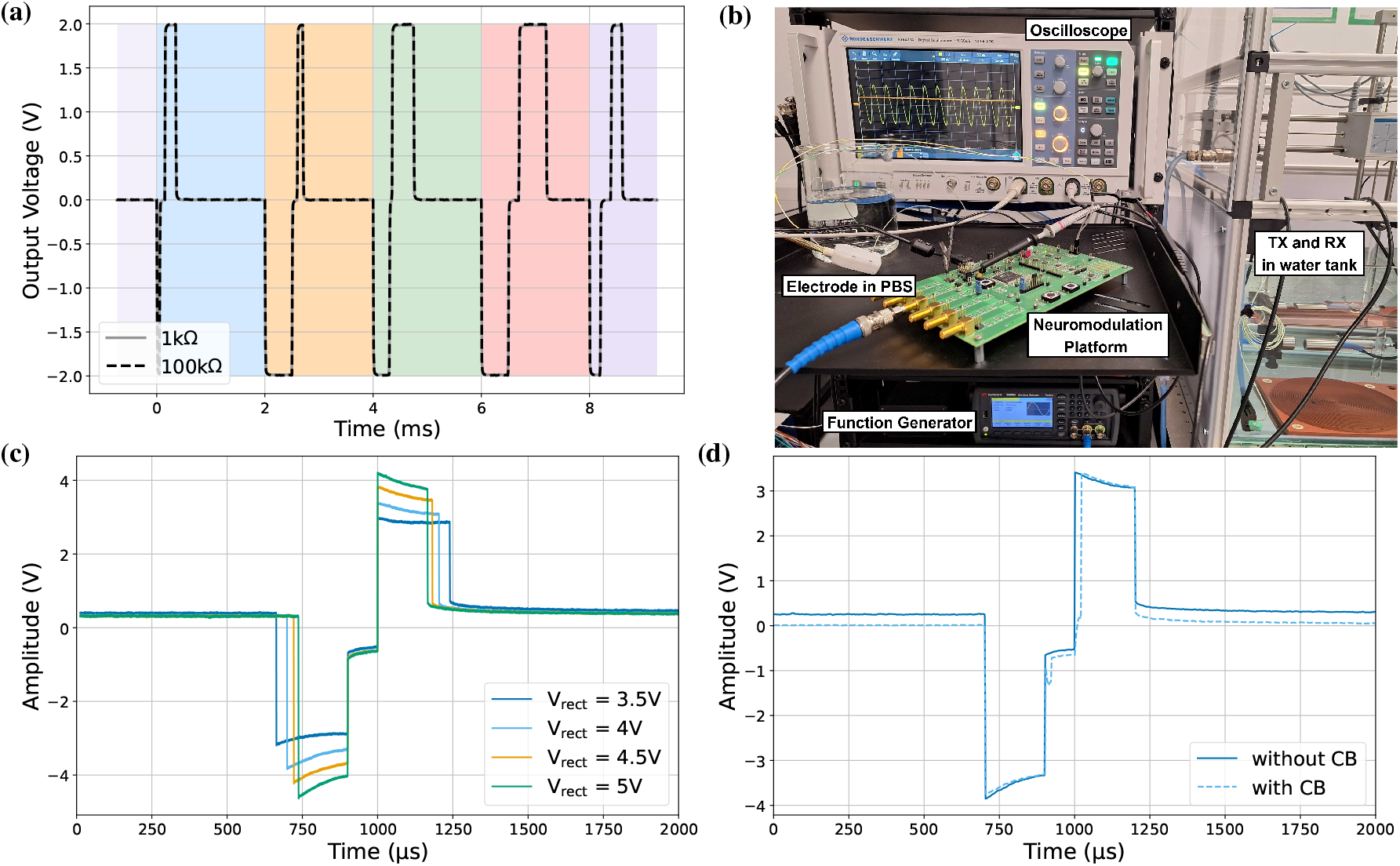
a) Measured voltage output of the neuromodulation platform, intended for neural activity excitation, when connected to a resistive load. Here, all five available patterns are activated, with the shaded areas corresponding to the colours of the respective pattern as displayed on the GUI. b) Neuromodulation platform characterization setup. The function generator drives the US transmitter TX, and the US receiver RX is connected to the platform for powering. Both the TX and RX are immersed in a water tank. The output of the platform is connected to the DRG leads from Abbott^©^, which are submerged in a PBS solution. The oscilloscope aids in visualizing the system’s output. c) Measured voltage output of the neuromodulation platform for a varying *V*_rect_ -the voltage used both for powering the platform and stimulating the electrodes connected to the output stage. The pulse width of each phase (cathodic and anodic) decreases/increases with increasing/decreasing *V*_rect_, respectively. d) Difference in the measured voltage output of the neuromodulation platform when having the charge balancing mechanism deactivated and subsequently activated. Here, we observe one biphasic pulse as part of a pulse sequence with a PRR of 100 Hz, and it is visible that the voltage offset between biphasic pulses is reduced significantly when activating the charge balancing (CB) mechanism.

#### G.2. Electrode Voltage Output

To evaluate the performance of the neuromodulation platform, the system is powered wirelessly through US, with the signal used for powering being forwarded to the output stage for stimulation. The output stage of the platform is connected to the DRG leads from Abbott^©^, submerged in a PBS solution to emulate the environment of the inner body. Part of the setup used for the system’s characterization is depicted in figure 8b.

Given that the voltage *V*_rect_ used for powering the platform is also used for stimulation, the stimulation pulses have an amplitude that is dependent on the intensity of the harvested US signal. In the event of transducer misalignment or movement, the harvested power might fluctuate, affecting the stimulation pulse amplitude. To ensure that the necessary amount of charge needed for neural stimulation is delivered within one stimulation pulse, the pulse width is scaled according to the varying *V*_rect_. Consequently, the pulse width of each stimulation phase will become longer for a lower *V*_rect_, or shorter for a higher *V*_rect_. Figure 8c demonstrates the implemented scaling at the example of four different voltages *V*_rect_.

Finally, the system’s output is observed when stimulating with the charge balancing mechanism deactivated and activated. At the example of one pulse within a stimulation sequence with a PRR of 100 Hz, the voltage offset between stimulation pulses decreases considerably while reducing the pulse width of the second (here, anodic) phase of the biphasic pulse. Figure 8d shows the measured output between two electrodes, without and with the charge balancing activated.

## Discussion

The presented platform offers a readily reproducible and versatile neuromodulation system that can be used to further explore the possibilities that VNS has to offer. Using only commercially available components, we developed a system that is intended for simultaneously exciting and inhibiting neural activity, while offering the possibility to connect it to multiple stimulating electrodes.

Powering the platform wirelessly through US opens the way for a future miniaturization and exploration of US-powered neuroelectronic implants that include only the most essential and power-efficient components needed for the intended application. Here, the wireless power transfer link consists of a commercially available US transmitter and US receiver, with an electrical impedance of 50 Ω. Though the platform allows for the incorporation of a matching network, this was not needed in this configuration. Working towards a miniaturized implant, the bulky commercial US receiver can be replaced by integrated piezoelectric or capacitive micromachined ultrasound transducers (PMUTs or CMUTs, respectively) (33, 34), which will require the introduction of a matching network.

The choice to use the same signal for powering and stimulation, despite improving the power efficiency of the system, comes at the cost of a stimulation pulse amplitude dependent on the US signal intensity. This dependency removes one degree of freedom for defining the stimulation parameters. It can, however, be compensated for by adjusting the width of the stimulation pulses with respect to the available power, to ensure that the necessary amount of charge needed for stimulation is delivered to the tissue. The corresponding scaling in this implementation is linear; it can, however, also follow a different trend, e.g., that of the strength-duration curve (35). Finally, the intensity of the received US signal is currently controlled through the definition of the sent US signal, ensuring that the resulting voltage is within the limits that the components of the platform can tolerate. The addition of a Zener diode in reverse bias can help to maintain this voltage within the required limits.

The integrated charge balancing technique increases stimulation safety by pursuing a zero net potential difference between all stimulating electrodes. For this, we define a reference against which we compare the potential of every electrode and place all electrodes (stimulating and reference) within the same environment. By attempting to bring every electrode to the potential of the reference electrode, the voltage offset between the electrodes is significantly reduced when enabling charge balancing, achieving offsets as low as 2 mV. Still, this is only a safety measure and does not provide a safety guarantee, as with most charge balancing mechanisms.

Notable is the accumulated voltage offset difference between pulses with lower versus higher PRRs, when the charge balancing mechanism is not yet activated. As can be seen in figure 7d, the potential difference between the stimulating electrodes is higher for lower PRRs; here at 300 mV for 100 Hz versus 200 mV for 1 kHz. This value, however, does not correspond to the typically measured voltage offset in between stimulation pulses of a pulse sequence, but rather to the mean voltage offset value over a longer period. This is because, to estimate the DC component that might lead to harmful electrochemical reactions at the ETI and eliminate it, we use the mean value of the full signal, including the stimulation pulses, as approximated by the incorporated LP filter. Therefore, even though a momentarily sampled voltage offset in between stimulation pulses would, as expected, measure lower for lower PRRs, the mean value over the full signal measures higher, given that this offset is present over a longer period when compared to a stimulation sequence with a higher PRR. By averaging the full signal, we take into consideration every instance of the stimulation pulse, including any hardware limitations (inconsistencies in the pulse widths, unwanted leakage currents, etc.) and the stimulation pulses themselves. The effectiveness of the chosen charge balancing mechanism is illustrated in 7d, lowering the voltage offset, independent of the PRR or the number of stimulation patterns, to a few mV.

Currently, the developed neuromodulation platform can be used as a benchtop or in an intraoperative setting, as it still requires a connection to a laptop or PC to set the stimulation parameters through the GUI. A future design will include wireless communication to decouple the device from any external connections completely, as it can already be powered through a battery or wirelessly through US (36).

## Conclusion

The neuromodulation platform presented in this work provides a practical and versatile experimental tool for investigating the functional potential of neuromodulation applications, including vagus nerve stimulation (VNS). Built around a commercially available microcontroller and off-the-shelf components, the system is wirelessly powered via ultrasound, leveraging the same signal for both energy transfer and stimulation to enhance efficiency. It supports both excitation and inhibition of neural activity, employing multiple electrodes for spatially distributed stimulation. To improve safety, the platform integrates a charge balancing mechanism, making it suitable for extended experimental use.

## ACKNOWLEDGEMENTS

This work was supported by the Moore4Medical project through the Electronics Components and Systems for European Leadership (ECSEL) Joint Undertaking under grant H2020-ECSEL-2019-IA-876190. This work was also funded as part of the Fraunhofer PREPARE Project “DUSTIN”. The authors would like to thank Andrada Iulia Velea for the practical help with the use of various types of ultrasound transducers.

